# Molecular Basis for Interferon-mediated Pathogen Restriction in Human Cells

**DOI:** 10.1101/2022.10.21.513197

**Authors:** Sumit K. Matta, Hinissan P. Kohio, Pallavi Chandra, Adam Brown, John G. Doench, Jennifer A. Philips, Siyuan Ding, L. David Sibley

## Abstract

To define novel mechanisms for cellular immunity to the intracellular pathogen *Toxoplasma gondii*, we performed a genome-wide CRISPR loss-of-function screen to provide an unbiased assessment of genes important for IFN-γ-dependent growth restriction. We revealed a previously unknown role for the tumor suppressor NF2/Merlin for maximum induction of Interferon Stimulated Genes (ISG), which are positively regulated by the transcription factor IRF-1. We then performed an additional focused ISG-targeted CRISPR screen that identified the host E3 ubiquitin ligase RNF213 as essential for IFN-γ mediated control of *T. gondii*. RNF213 mediated ubiquitination of targets on the parasite-containing vacuole and growth restriction in response to IFN-γ in a variety of cell types, thus identifying a conserved factor that plays a prominent role in human cells. Surprisingly, growth inhibition did not require the autophagy protein ATG5, indicating that RNF213 initiates restriction independent of a non-canonical autophagy pathway that has previously been implicated in control of *T. gondii*. RNF213 was also important for control of unrelated intracellular pathogens in human cells treated with IFN, as shown here for *Mycobacterium tuberculosis* and Vesicular Stomatitis Virus. Collectively, our findings establish RNF213 as a critical component of cell-autonomous immunity to a broad spectrum of intracellular pathogens in human cells.

## Introduction

*Toxoplasma gondii* is a widespread parasite of animals that also frequently causes zoonotic infection in humans. As an obligate intracellular parasite, *T. gondii* resides within a specialized vacuole that is largely sequestered from the host endomembrane system (1), which the parasite protects to provide niche for intracellular growth (2, 3). Cell autonomous mechanisms for immune control of intracellular *T. gondii* are best known from studies in the natural murine host, for which laboratory rodents are a useful model (4, 5). Type II interferon, or interferon gamma (IFN-γ), plays the major role in resistance to infection in mouse (6, 7), while type I interferon is also produced and contributes to control of infection (8), particularly in the central nervous system (CNS) during chronic infection (9). Treatment with IFNs leads to upregulation of numerous Interferon Stimulated Genes (ISGs), the exact complement of which varies by cell type (10, 11). Among the most strongly upregulated ISGs are the critically important immunity related GTPases (IRGs) and guanylate binding proteins (GBPs), which collectively lead to vacuole rupture, parasite destruction, and clearance (12-14). The IRG/GBP system is under control of a noncanonical autophagy (ATG) system that requires core ATG components for conjugation of LC3 to phosphatidyl ethanolamine (PE), but it does not require the initiation nor degradative steps (15-17). Although, IRGs and GBPs play a central role in immunity in rodents, human cells largely lack IRGs and GBPs have been described to play a more limited role, as discussed further below.

Clinical evidence that human cells control intracellular *T. gondii* via an interferon-mediated process comes from studies demonstrating that immunocompromised patients with low levels of IFN-γ are susceptible to reactivation of latent toxoplasmosis (18, 19). In vitro studies indicate that multiple IFN-γ-dependent pathways can mediate growth restriction in humans. Although humans are not natural hosts in the life cycle, human macrophages are nonetheless able to control the replication of intracellular *T. gondii* in response to IFN-γ (20, 21) and IFN-γ (22, 23). Nonhematopoietic human cells also control intracellular *T. gondii* through varied and often cell-type specific mechanisms (24). For example, induction of indole amine oxidase (IDO) by interferon leads to degradation of tryptophan, thus limiting parasite growth in human fibroblasts (25) and human brain microvascular endothelial cells (26). In addition, the noncanonical autophagy pathway described above is required for recruitment of LC3, leading to engulfment of the parasite containing vacuole in multiple membranes, a process that is associated with growth restriction in Hela cells (27), A549 cells (28), and human umbilical vein endothelial cells (29). Curiously, the ATG-dependent growth restriction is not global (16), but rather limited to those vacuoles that become ubiquitinated and acquire adaptors such as p62, NDP52, and LC3 (27, 29). Virulent type I strains are largely resistant to this pathway, while types II and III are susceptible (27), although the basis for this difference is unknown. Recent studies identified the E3 ubiquitin ligase RNF213 as a candidate for mediating early ubiquitination of *T. gondii-* containing vacuoles in IFN-γ treated A549 cells (30). The IFN-γ signaling and noncanonical ATG pathways are partially connected by ISG15, which is required for maximal recruitment of ATG adaptors and also interacts with RNF213 (28). Additionally, GBP1 has been implicated in restricting *T. gondii* growth in human epithelial cells, despite not being recruited to the vacuole (31), while it is recruited to the vacuole membrane in THP-1 cells, resulting in rupture and activation of apoptosis (32). Additionally GBP2 and GBP5 contribute to growth restriction in THP-1 cells, despite not being recruited to the parasite containing vacuole (33). Collectively, these prior candidate-gene studies indicate that multiple IFN-γ–dependent mechanisms can contribute to growth restriction, which is often partial and expressed in a cell-specific manner. However, the absence of any prior global or unbiased assessment of genes required for IFN-γ mediated growth restriction leaves open the possibility that a previously unrecognized, IFN-γ-mediated pathway exists in all or most human cells.

To provide an unbiased assessment of the predominate pathways for control of *T. gondii* growth in IFN-γ activated human cells, we performed genome-wide CRISPR screen in A549 cells. Our findings reveal a previously unanticipated role for the tumor suppressor NF2 in modulating IRF-1-dependent gene expression, hence dampening ISG expression. Additionally, an ISG-focused CRISPR screen identified the E3 ligase RNF213 as an essential component of control of *T. gondii* in multiple human cell lines treated with IFN-γ. Surprising, although RNF213 was associated with ubiquitination and recruitment of ATG adaptors including LC3, it functioned independently of the previously described pathway for non-canonical autophagy. Moreover, RNF213 exerted a general antimicrobial activity against intracellular bacteria and viruses, including *Mycobacterium tuberculosis* and vesicular stomatitis virus, revealing a conserved role in cell autonomous immunity in human cells.

## Results

### Genome-wide CRISPR screen identifies factors important for IFN-γ mediated growth restriction

Although depletion of tryptophan due to induction of IDO has been described as a mechanism for restricting growth of *T. gondii* in some cell types (24), we previously reported that disruption of the IDO gene in A549 cells has no impact on IFN-γ mediated growth restriction (34). Consequently, we selected A549 cells to define novel mechanisms for IFN-γ mediate growth restriction based on CRISPR-mediated loss of function screens. Initially, we validated the growth restriction following IFN-γ activation of A549 cells using a previously described vacuole size assay (9). We used the type III strain named CTG of *T. gondii* as it is highly susceptible to growth restriction, relative to either type I or type II strains (27, 35). The vacuole size assay takes advantage of the fact that parasite replication leads to increased size of the vacuole while growth inhibition by IFN-γ blunts the expansion of parasites. Quantification of vacuole size after IFN-γ treatment verified that wild type (WT) cells restricted parasite expansion and that this ability was IFN-γ dependent, as shown by loss of growth restriction in STAT1 knockout (STAT1-KO) cells (**Fig. 1A, Fig. S1A**). As expected, the growth restriction was independent of exogenously added tryptophan, indicating it does not rely on IDO1, even though this gene is induced in IFN-γ treated A549 cells (34). To develop a process for selecting cells from the population, we infected A549 cells with a recombinant CTG strain expressing GFP (CTG-GFP), trypsinized cells from the dish, and analyzed them by flow cytometry to evaluate the average intensity of GFP as a proxy for parasite growth (**Fig. 1B**). Although there was not a distinct separation between the populations of CTG-GFP in WT vs. STAT1-KO cells, there was a clear shoulder of higher GFP expression in the STAT1-KO cells (green box, **Fig. 1B**). We therefore reasoned that escape mutants that lose the ability to control parasite replication should mimic the growth of CTG-GFP in STAT1-KO cells, thus providing the basis for the CRISPR-based screens described below.

**Figure 1.**
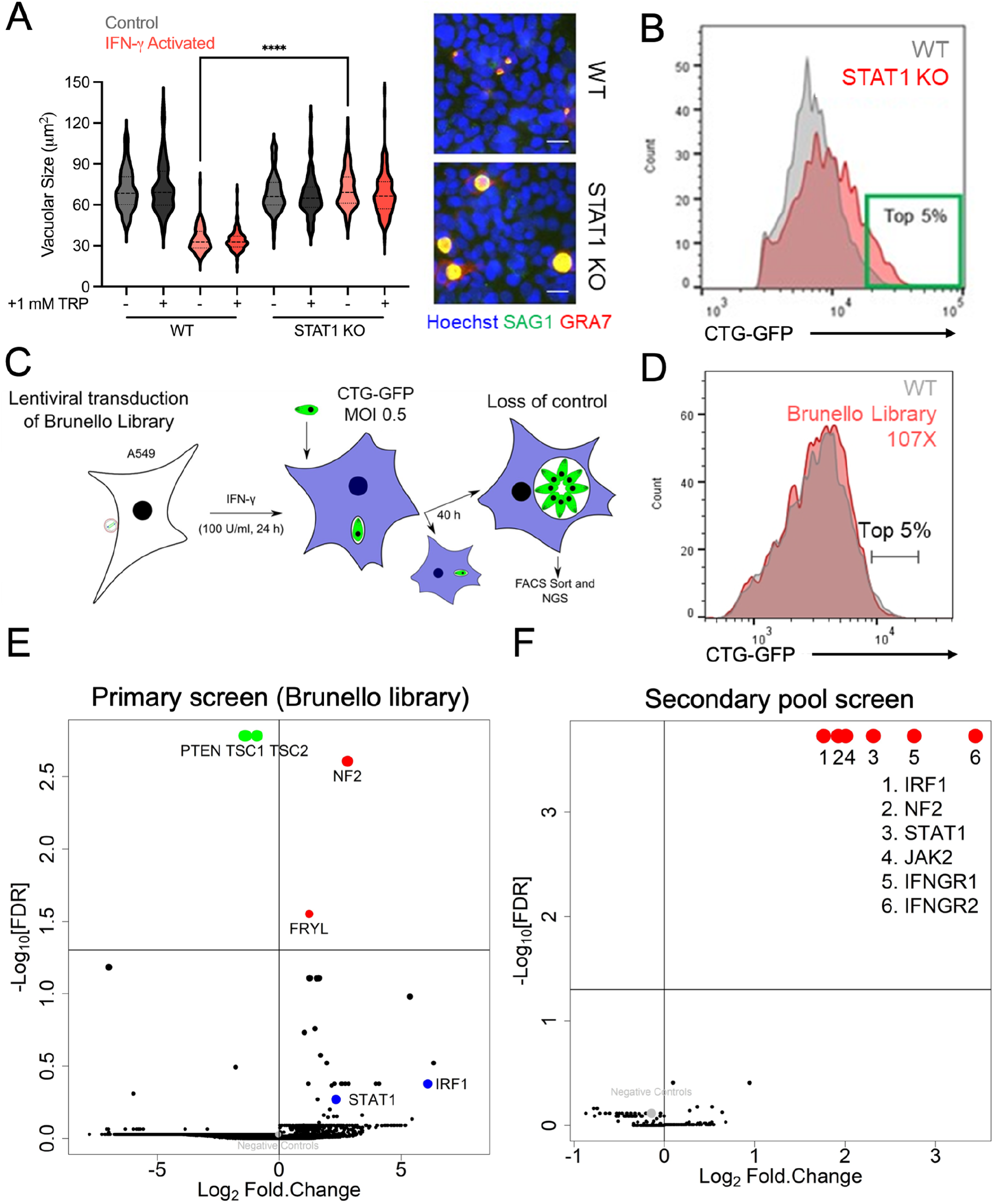
Design and implementation of genome-wide CRISPR screens for loss of IFN-γ-mediated parasite growth restriction in human cells (A) Tryptophan (TRP)-independent restriction of *T. gondii* in lung alveolar epithelial A549 cells. Wild type (WT) and STAT1 knock out (KO) A549 cells (**Fig. S1A**) were treated ± IFN-γ (100 Units/ml) 24 h prior to infection with type III CTG strain of *T. gondii*. The cells were either left untreated (Control) or supplemented with 1 mM TRP for 40 h. Growth was measured by vacuolar size (microns^2^) for at least 3 biological replicates with ≥ 30 images per sample and replicate. Violin plot shows mean vacuolar size per image with black bar representing the median. **** *P* < 0.0001 using unpaired Two-tailed t-test with Welch’s correction. Representative images of WT and STAT1 KO A549 cells (+IFN-γ +TRP) infected with CTG and analyzed at 40 h hpi. Nuclei labeled with Hoechst (blue), parasites with anti-SAG1 (green) and the vacuole membrane with anti-GRA7 (red). Scale bar = 20 microns. (B) IFN-γ activated WT vs. STAT1 KO A549 cells infected with CTG expressing GFP (CTG-GFP) were fixed 40 hpi and analyzed by flow cytometry. Histogram shows GFP positive (CTG-GFP infected) cells. Green box defines the top 5% of CTG-GFP expressing cells. (C) Design of genome-wide CRISPR-Cas9 to identify factors involved in IFN-γ mediated parasite growth restriction in human (A549) cells. (D) The Brunello library (36) was transduced into A549 cells at ∼100X coverage. Histogram of CTG-GFP infection analyzed at 40 hpi in IFN-γ activated control (WT) and Brunello library transduced A549 cells. (E) Primary genome-wide CRISPR screen showing Log_2_ fold change in sgRNAs from the top 5% of CTG-GFP expressing cells (T5) compared to uninfected cells plotted against -log_10_ false discovery rate (FDR) for all 4 independent replicates combined. Genes for significantly enriched sgRNAs (red) vs. significantly depleted sgRNAs in the T5 pool (green) are labeled. (F) Secondary CRISPR screen consisting of a sub-pool of 224 genes that were significantly enriched in the T5 population (Log_2_Fold Change > 1 and *P* < 0.05) in at least 2 out of the 4 replicates of the primary Brunello screen (**Fig. S1B**). Log_2_ Fold Change is plotted against -log_10_ FDR for all 4 independent replicates combined.

We then used lentiviral transduction of A549 cells to express the Brunello library containing 4 guides each to ∼20,000 genes in the human genome (36) and developed a screen for loss of IFN-γ mediate growth restriction (**Fig. 1C**). We scaled the culture assays to aim for single sgRNA per cell with ∼ 100x coverage of each gene such that loss of guides during expansion could be quantified. Cells were expanded for 8 passages to allow for stabilization of the population of guides that did not affect viability. Cells were then activated with IFN-γ, infected with CTG-GFP, and the top 5% of GFP expressing cells were sorted from four independent experiments (**Fig. 1D**). The sgRNAs from these pools were amplified, barcoded, and subjected to NGS sequencing using the Illumina platform. We reasoned that sgRNAs that are enriched in the top 5% pool represent genes whose loss results in impaired IFN-γ mediated growth restriction of CTG-GFP. Comparison of the guide abundances in the top 5% relative to a time-matched point for the uninfected pool was used to classify guides that were significantly different based on Fold Change and FDR (False Discovery Rate) in the four replicates of the primary screen (**Fig. 1E, Dataset 1**). There was a high degree of variability in genes that were scored significant between the various replicates. For example, only 5 genes (including NF2, FRYL) met the individual *P* value for significance in all 4 replicates when analyzed separately (**Fig. S1B**). Additionally, 23 genes were individually significant in 3 of 4 while 224 were significant in 2 of 4 replicates (**Fig. S1B**). However, using the more stringent criteria of FDR for the combined replicates, only NF2 and FRYL were scored as significant (**Fig. 1E**). Although guides for STAT1 and IRF1 were slightly enriched, the strongest signal for enhanced sgRNA guides correspond the gene NF2 (Neurofibromatosis 2) an Ezrin / Radixin / Moesin (ERM) family member that is also known as MERLIN (37), and which functions as a tumor suppressor (**Fig. 1E**). We also detected three genes that showed depletion of sgRNAs (PTEN, TSC1 and TSC2) (**Fig. 1E**). Although we have not followed up on these hits, they presumably are depleted because cells lacking these genes have decreased survival during infection and IFN-γ treatment.

### Secondary pool CRISPR screen identifies IFN-γ-STAT1 axis and NF2 as critical for growth restriction

Comparison of the results from the primary screen revealed a high degree of variability in genes classified as significantly different between the replicates, which is not unexpected for FACS-based screens (38). We therefore considered it possible that the genome-wide screen may have missed genes that contribute to IFN-γ mediated resistance. Consequently, we developed a secondary screen based on a pool of genes that were significantly enriched in at least 2 of 4 replicates of the primary screen (**Fig. S1B**). The secondary screen included 224 genes with coverage of 10 sgRNAs per gene, thus improving statistical power for identifying differences. The secondary screen identified several factors in the IFN-γ-STAT1 axis; their loss suggests that they are essential for growth restriction **(Fig. 1F**) (**Dataset 2**). Included in the set of highly enriched guides representing depleted genes were STAT1, the JAK2 kinase that phosphorylates STAT1, the IFN-γ receptor subunits IFNGR1 and IFNGR2 as well as the downstream transcription factor IRF-1, which augments the activity of IFN-γ on inducing gene expression (39) (**Fig. 1F**). The only gene not part of the IFN-γ-STAT1 pathway was NF2, which was the top hit in the primary screen (**Fig. 1F**). The four other genes that were significantly enriched in each replicate of primary screen, including FRYL, were not enriched in the T5 population of the secondary screen.

NF2 has previously been implicated in suppressing tumors via interaction with the E3 ubiquitin ligase CRL4 in the nucleus (37), and in regulating mTOR (40) and modulating metabolism (41); however, it has not previously been associated with IFN-γ activation nor pathogen control. To elucidate its contribution to cell-autonomous pathogen control, we generated KOs using lentiviral transduction and targeted CRISPR-Cas9 gene editing for both NF2 and IRF-1 in A549 cells (**Fig. S2**). As expected, IRF-1 showed dependence for expression in WT cells and NF2-KO cells on IFN-γ treatment, while NF2 was not affected by IFN-γ treatment (**Fig. 2A**). We then ascertained the ability of IFN-γ to control growth of CTG parasites in WT, IRF-1 KO and NF2 KO cells using the vacuole size assay. As expected, growth restriction seen in WT cells was completely lost in IRF-1 KO cells, consistent with previous reports that this transcription factor is essential for control of *T. gondii* in IFN-γ treated cells (34). In contrast, loss of NF2 resulted in a partial, yet significant reduction in the growth restriction following IFN-γ (**Fig. 2B and 2C**).

**Figure 2.**
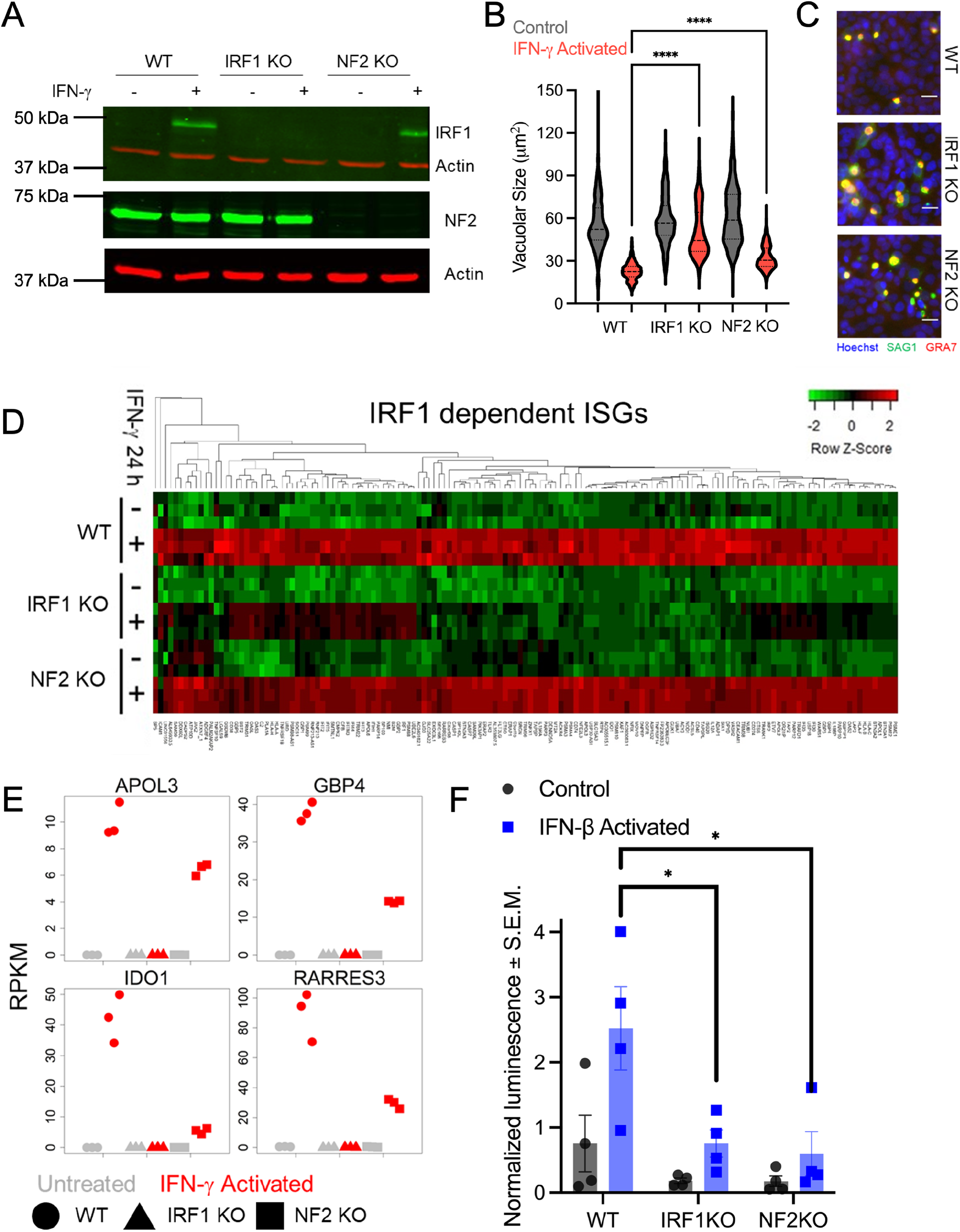
NF2 dampens IFN-γ-mediated growth restriction by inhibiting ISG induction. (A) Immunoblots for IRF-1 and NF2 in wild type, IRF1 KO, and NF2 KO A549 cells (**Fig. S2A and S2B**). Actin was used as a loading control. (B) Wild type, IRF1 KO, and NF2 KO A549 cells treated ± IFN-γ (100 U/ml) for 24 h, infected with CTG strain *T. gondii*, and evaluated for growth restriction at 40 hpi. Violin plots show mean vacuolar size per image with the black bar representing the median for at least 3 biological replicates containing at least 30 images per sample and replicate. **** *P* < 0.0001 using Ordinary one-way ANOVA with Turkey’s multiple comparison test. (C) Representative images of IFN-γ activated wild type, IRF1 KO and NF2 KO A549 cells infected with CTG and analyzed at 40 hpi. Cells were labelled with Hoechst for nuclei (blue), anti-SAG1 for parasites (green), and anti-GRA7 for vacuoles (red). Scale bar = 20 microns. (D) Global transcriptional profiling from A549 cells treated ± IFN-γ (100 U/ml) for 24 h. Heat maps depict genes that were induced in WT but not in IRF1 KO cells from 3 biological replicates clustered using 1-Pearson distance and average linkage on normalized Log_2_ (RPKM) values. Normalized Z-scores are color-scaled from down-regulation (green) to upregulation (red). (E) RPKM values showing partial induction of select ISGs in NF2 KO cells upon IFN-γ activation compared to WT cells. (F) ISRE promoter activity from A549 cells transfected with ISRE reporter plasmid expressing Gaussia luciferase (9) and control pCMV-Red Firefly Luc plasmid (Thermo Fisher Scientific) for 24 h. Bar plot shows mean ± S.E.M Gaussia luciferase activity normalized to firefly luciferase activity for 4 independent biological replicates. * *P* < 0.05 using Two-way ANOVA with Turkey’s multiple comparison test.

### Loss of NF2 down-modulates IRF-1 dependent ISG expression

To better define the role of NF2 in IFN-γ dependent growth restriction, we compared the transcriptional profile of WT, IRF-1 KO and NF2 KO cells under resting and activated conditions. Comparison of the NF2 KO to WT cells revealed multiple of genes that were NF2 dependent (**Fig. S3**), which likely reflect its diverse roles. However, when we compared the IFN-γ induced genes that were IRF-1-dependent, there was a clear pattern that most of these genes were also dampened in the NF2 KO (**Fig. 2D**). The extent of suppression in the NF2 KO varied among genes: for example, APOL3 was IRF-1 dependent but only partially affected by NF2, whereas IDO, GBP4 and RARESS3 were more strongly affected by NF2 as well as IRF-1 (**Fig. 2E**). We also examined the effect of NF2 on IFN-β driven expression using an ISRE (Interferon Stimulation Response Element) promoter normalized to constitutively expressed firefly luciferase. Induction of ISRE activity by IFN-β was significantly lost in both IRF1 KO and NF2 KO cells (**Fig. 2F**), indicating that both factors likely also participate in IRF-1 mediated transcription.

### ISG targeted CRISPR screen identifies RNF213 as the primary determinant of IFN-γ mediated growth restriction in multiple human cell types

We were surprised that neither the primary nor secondary genome-wide screens identified any canonical ISGs, except for the broadly acting transcription factor IRF-1, which affects many ISGs. Consequently, we designed a focused CRISPR screen based on a previous set of ISGs that was defined by transcriptional induction in response to IFN-γ in A549 cells (34). The ISG screen included ∼350 genes with coverage at 10 sgRNAs per gene and was repeated four times independently (**Fig. 3A**). Sequencing the top 5% of CTG-GFP positive cells was used to define sgRNAs that were significantly enriched, thus identifying genes important for IFN-γ mediated control. We identified only a single strong positive corresponding the E3 ligase RNF213, in addition to the broadly acting IRF-1 (**Fig. 3B**) (**Dataset 3**). RNF213 is not widely recognized as an ISG due to its constitutive expression in many cell types, and yet it is upregulated by ∼ 8 fold in A549 cells treated with IFN-γ and thus was part of the library used in the screen. RNF213 ranked 304 in the primary screen (**Dataset 1**) but did not meet the criteria for the secondary pool screen, although sgRNA guides for this gene were enriched in the top 5% population in 3 of 4 replicates by > 2 fold. Given its prominent position in the targeted ISG screen, we decided to examine its function. Deletion of RNF213 resulted in enhanced growth of CTG strain parasite in both untreated cells and it completely reversed the growth inhibition normally seen in IFN-γ treated A549 cells (**Fig. 3C and 3D**). Notably, although sgRNAs were present for IDO1, ISG15, GBP1, and RARRES3, their frequency was not changed in the top 5% pool, indicating the singular loss of these genes does not contribute substantially to IFN-γ mediated growth restriction under the conditions of the screen.

**Figure 3.**
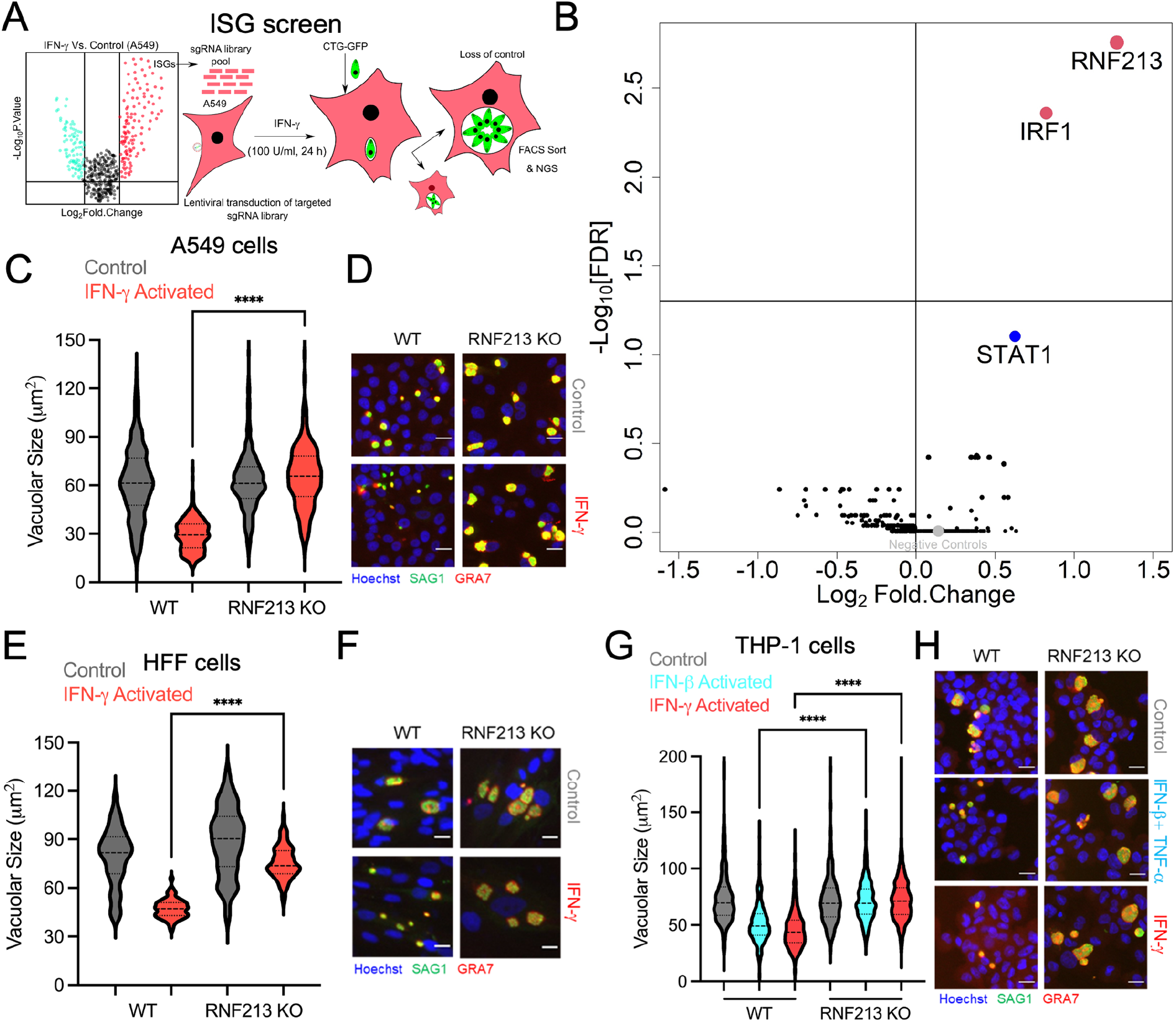
CRISPR-Cas9 screen identifies RNF213 as the primary ISG required for IFN-γ-mediated growth restriction of *T. gondii* in human cells (A) Design of ISG targeted sgRNA CRISPR-Cas9 screen using a sub-pool of ISGs that were previously shown to be upregulated in A549 cells treated with IFN-γ (34). (B) Targeted sgRNA screen of ISGs upregulated by IFN-γ identified two genes that were significantly depleted in the top 5% of CTG-GFP expressing cells (red). Log_2_ Fold Change is plotted against -log_10_ FDR for all 4 independent replicates combined. (C) Wild type and RNF213 KO A549 cells (**Fig. S4A and 4B**) treated ± IFN-g (100 U/ml) for 24 h, infected with CTG strain *T. gondii*, and evaluated for growth restriction by quantifying vacuolar size. Vacuolar growth of parasites was measured for at least 3 independent biological replicates with 30 images per sample and replicate. Violin plot shows mean vacuolar size per image with black bar representing the median. \*\*\*\**P* < 0.01 using two-tailed unpaired Student’s t-test with Welch’s correction. (D) Representative images showing nuclei stained with Hoechst (blue), parasites stained with anit-SAG1 (green), and the vacuole membrane stained with anti-GRA7 (red). Scale bar = 20 microns. (E) Vacuolar growth of parasites at 40 hpi in ± IFN-γ activated WT and RNF213 KO human foreskin fibroblast (HFF) cells. Violin plot shows mean vacuolar size (μ m^2^) per image with black bar as median for at least 3 independent biological replicates with at least 30 images per sample and replicate. \*\**\*\* P* < 0.0001 two-tailed unpaired Student’s t-test with Welch’s correction. (F) Representative images of the vacuole size in control and IFN-γ activated WT and RNF213 KO HFF cells infected with CTG and analyzed at 40 hpi. Nuclei were labelled with Hoechst (blue), the parasite with anti-SAG1 (green) and the vacuole membrane with anti-GRA7 (red) with bar = 20 μ m. (G) Vacuolar growth of parasites at 40 hpi in control, IFN-γ +TNF-α and IFN-γ activated WT and RNF213 KO human THP-1 macrophages. Violin plot shows mean vacuolar size (μ m^2^) per image with black bar as median for at least 3 biological replicates with at least 30 images per sample and replicate. \*\**\*\* P* < 0.0001 using Ordinary one-way ANOVA with Turkey’s multiple comparisons test. (H) Representative images of control, IFN-γ +TNF-α and IFN-γ activated WT and RNF213 KO THP-1 cells infected with CTG and analyzed at 40 hpi. Nuclei are labelled with Hoechst (blue), parasites with anti-SAG1 (green) and the vacuole membrane with anti-GRA7 (red). Scale bar = 20 μm.

Many factors that have been previously described to restrict *T. gondii* growth in IFN-γ treated human cells are either cell type specific or limited to specific parasite lineages (24). To test how widely conserved the function of RNF213 was, we generated KOs in human foreskin fibroblasts (HFF) and in the human monocytic cell line THP-1 (**Fig. S4**). Similar to the situation in A549 cells, loss of RNF213 resulted in slightly enhanced growth in untreated HFF cells and a loss of IFN-γ mediated growth inhibition (**Fig. 3E and F**). Additionally, the loss of RNF213 in THP-1 cells reversed the inhibition of *T. gondii* growth following treatment with either IFN-γ or IFN-β combined with TNF-α (**Fig. 3G and H**), either of which have previously been shown to control *T. gondii* growth (9). Notably, the loss of RNF213 in all three cell types completely reversed the effects of IFN-γ, similar to loss of STAT1 or IRF-1 in A549 cells. Additionally, the expression of RNF213 was sub-maximal in IRF-1 KO and NF2 KO cells treated with IFN-γ (**Fig. S4A**), suggesting they may act through this downstream mediator. Collectively, these findings argue that RNF213 is the major effector downstream of both type I and type II IFNs that inhibits parasite growth in human cells.

### RNF213-dependent growth restriction leads to recruitment of autophagy adaptors but is independent of ATG5

Consistent with its requirement for growth restriction, RNF213 was recruited to a portion of *T. gondii* vacuoles in untreated cells, and this increased dramatically following treatment with IFN-γ as shown by immunofluorescence assay (IFA) and immuno-EM (**Fig. 4A-C**). Since ubiquitination is thought to be the first step in marking the *T. gondii* vacuole for growth restriction (27), we stained cells with FK2, which recognizes both mono and poly ubiquitin (Ub) chains, in combination with RNF213. The proportion of vacuoles that were FK2 positive closely matched the percentage that were also positive for RNF213, while the number that were RNF213 positive considerably exceeded this value (**Fig. 4C**). The higher percentage of RNF213 positive staining vacuoles, suggesting that RNF213 is initially recruited to the PVM followed by ubiquitination, resulting in double positive vacuoles. Similar co-staining of RNF213 and FK2 was observed in IFN-γ activated HFF and THP-1 cells infected with CTG (**Fig. S5)**. Downstream of IFN-γ mediated ubiquitination, the PVM acquires several adaptors, including p62 and NDP52, that recruit the autophagy adaptor LC3 (27-29). Consistent with these previous reports, recruitment of LC3 was low in non-activated cells, and significantly elevated after treatment with IFN-γ (**Fig. 4D and E**). The proportion of vacuoles that were positive for both FK2 and LC3 was similar, while singly FK2 positive vacuoles considerably exceeded this value (**Fig. 4E**). These differences in positivity suggests that ubiquitination occurs first, followed by recruitment of LC3.

**Figure 4.**
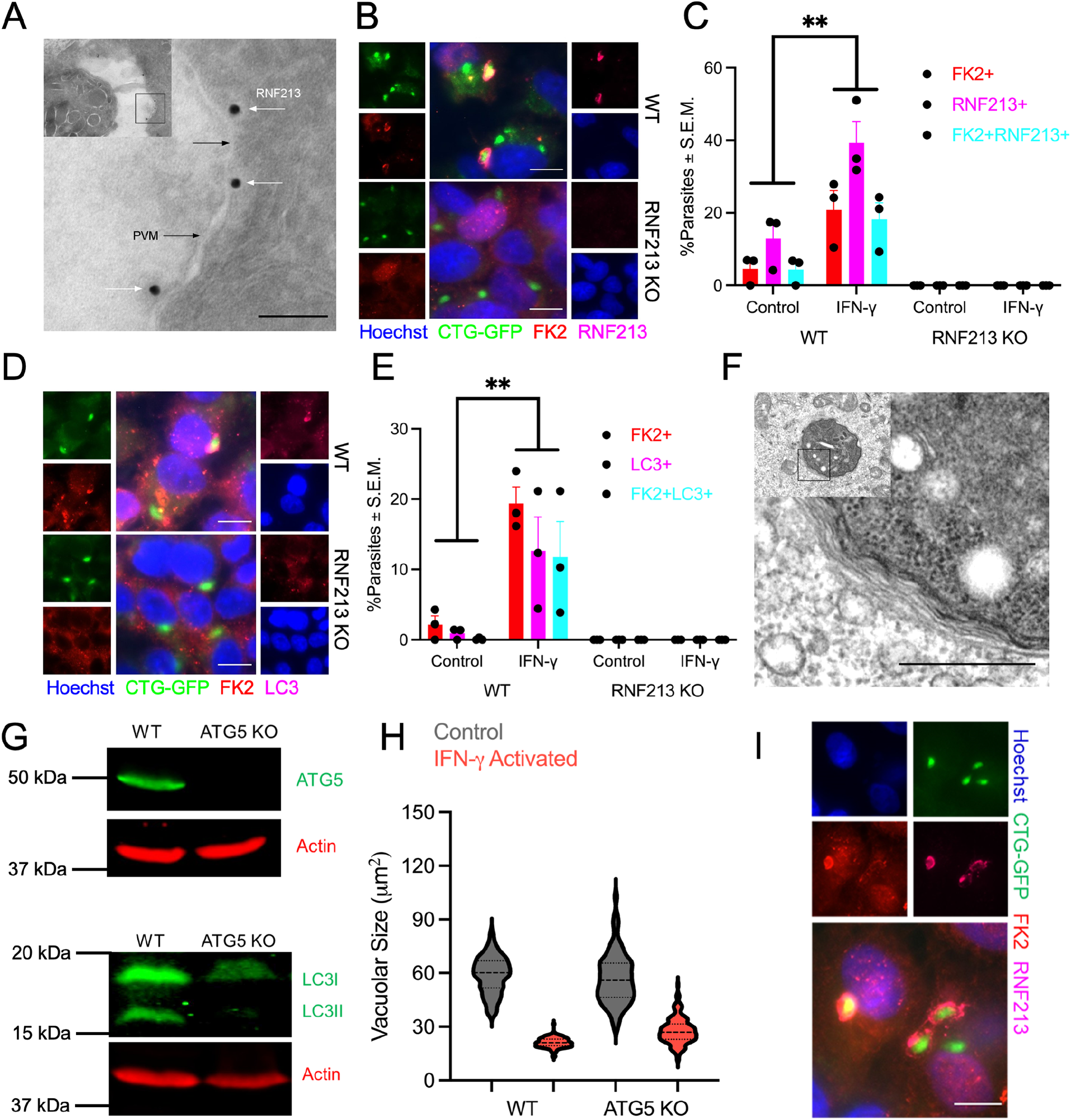
RNF213 co-localized with ubiquitinated parasite-containing vacuoles. (A) Immuno-EM labeling of RNF213 in IFN-γ activated wild type A549 cell infected with CTG parasite (inset) at 6 hpi. Parasite vacuolar membrane (PVM). Bar = 100 nm. (B) Representative images of WT and RNF213 KO A549 cells infected with CTG-GFP at 6 hpi. Cells were stained with FK2 (mono-poly ubiquitin; red) and anti-RNF213 (magenta). Nuclei were labelled with Hoechst (blue) and parasites were GFP labelled (green), Bar = 10 μm. (C) Quantification of FK2+ (red), RNF213+ (magenta) and FK2+RNF213+ (cyan) parasites for at least 3 biological replicates, mean ± S.E.M. ** *P* < 0.01 using Two-way ANOVA with Turkey’s multiple comparisons test. (D) Representative images of wild type and RNF213 KO A549 cells infected with CTG-GFP at 6 hpi. The cells were stained with FK2 (red) and anti-LC3B (magenta). Nuclei were labelled with Hoechst (blue) and parasites were GFP labelled (green). Scale bar = 10 μm. (E) Quantification of FK2+ (red), LC3+ (magenta) and FK2+LC3+ (cyan) parasites for at least 3 biological replicates, mean ± S.E.M. ** *P* < 0.01 using Two-way ANOVA with Turkey’s multiple comparisons test. (F) Transmission electron microscopic image of IFN-γ activated wild type A549 cell infected with CTG parasite (inset) at 6 hpi. Wrapping by multiple membranes is highlighted in the magnified view. Bar = 500 nm. (G) ATG5 (top) and LC3B (bottom) immunoblots of WT and ATG5 KO A549 cells. Actin was used as loading control. (H) Vacuolar growth of CTG parasite at 40 hpi in ± IFN-γ activated wild type and ATG KO cells. Violin plot shows mean vacuolar size (μm^2^) per image with black bar as median for an experiment with at least 30 images per sample. (I) Representative image of IFN-γ activated ATG5 KO A549 cells infected with CTG-GFP at 6 hpi. The cells are stained with FK2 (red) and anti-RNF213 (magenta). Bar = 10 μm.

Examination of the vacuole membrane surrounding the parasite by TEM revealed that it was often wrapped by numerous membranes (**Fig. 4F**), a phenotype that has previously been associated with growth restriction by non-canonical autophagy (27). Combined with the recruitment LC3, this profile suggested that RNF213 may act to recruit components of the non-canonical autophagy pathway to restriction growth of the parasite. To test this idea, we generated a knockout of ATG5 in A549 cells, and verified its absence by western blotting and loss of conversion of LC3I to the lipidated LC3II form (**Fig. 4G**). Contrary to expectation, loss of ATG5 minimally affected the growth restriction of *T. gondii* in IFN-γ treated cells (**Fig. 4H**). To ascertain whether loss of ATG5 might have affected the recruitment of RNF213 to the parasite containing vacuole, we stained IFN-γ treated and infected ATG5 KO cells for RNF213 and FK2. Similar to WT cells, parasites were frequently labeled by both RNF213 and FK2 in ATG5 KO cells (**Fig. 4I**). Collectively, these findings indicate that RNF213 lies upstream of ubiquitination and that growth restriction requires RNF213 but not an ATG5 dependent process.

### RNF213 participates in interferon-mediated control of bacterial and viral pathogens

Previous studies have described a role for RNF213 in recognition and ubiquitination of LPS on intracellular *Salmonella* (42). To explore the role of RNF213 in control of intracellular bacteria that do not contain LPS, we examined the requirement for RNF213 in IFN-γ activated A549 cells infected with *Mycobacterium tuberculosis* (Mtb). Human cells normally are not able to clear intracellular Mtb following treatment with IFN-γ but rather restrict their intracellular growth. Consistent with this profile, the colony forming units of Mtb remained static in IFN-γ treated cells grown for 48 h, compared to control cells where they increased significantly (**Fig. 5A**). Loss of RNF213 significantly impaired the ability of IFN-γ to control replication of Mtb (**Fig. 5A**). Immunofluorescence localization also revealed that RNF213 was recruited to the surface of Mtb in A549 cells and that these vacuoles also stained with FK2 indicating that they were ubiquitinated (**Fig. 5B**). RNF213 has also been reported to facilitate control of Rift Valley fever virus, an enveloped segmented RNA virus (43). We also explored the potentially broad anti-microbial role of RNF213 in control of intracellular viral replication using a non-segmented RNA virus. We examined type I IFN (IFN-γ) treated A549 cells infected with vesicular stomatitis virus expressing GFP (VSV-GFP). Treatment with IFN-β resulted in a significant decrease in viral replication in WT cells that was reversed in IRF-1 KO and RNF213 KO cells (**Fig. 5C and D**). Collectively, these findings suggest that RNF213 may recognize a host component that is common to multiple pathogens, rather than a panoply of different molecules each unique to one pathogen.

**Figure 5.**
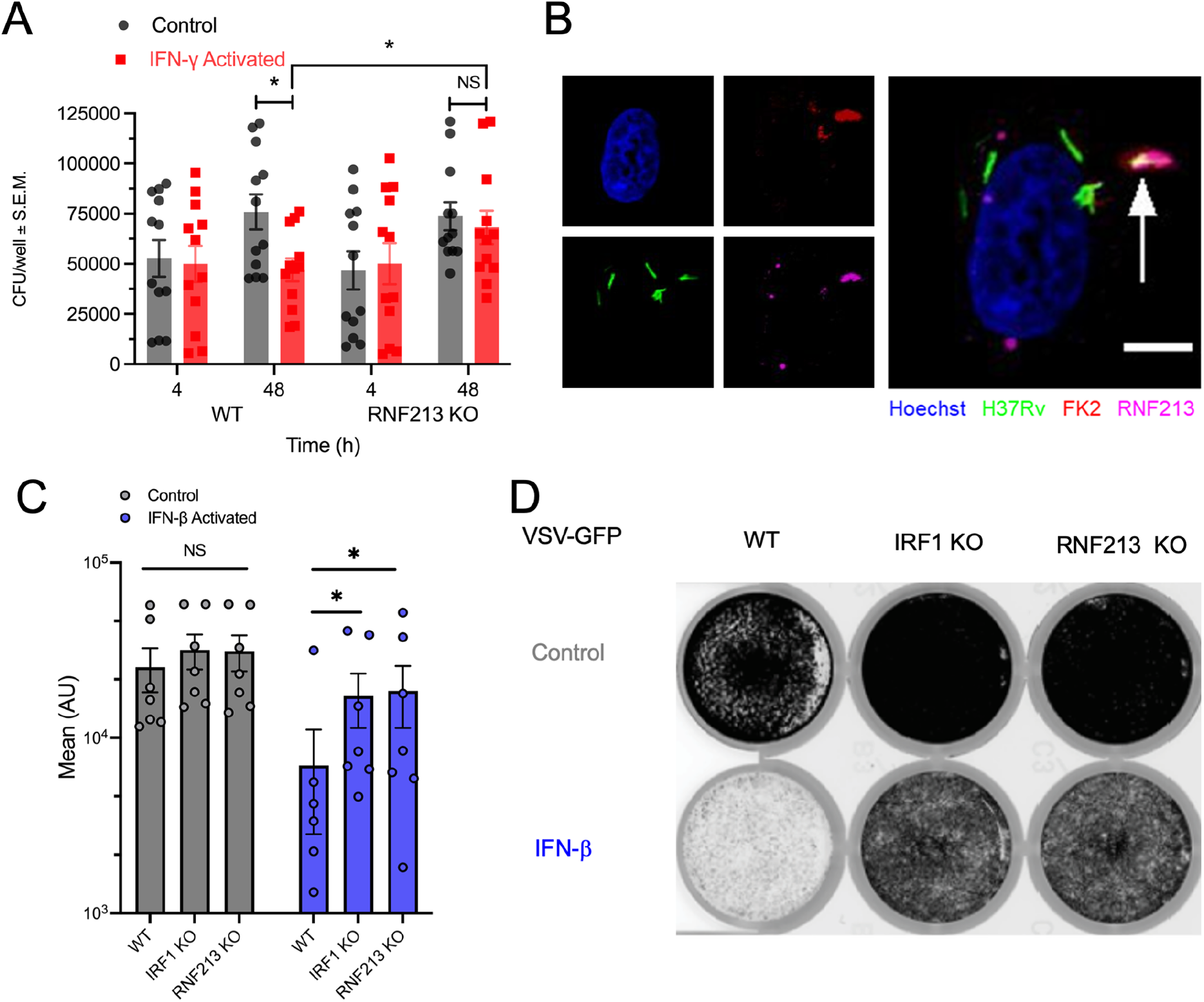
RNF213 is required for interferon-mediated control of intracellular bacterial and viral infection in human cells. (A) Mtb growth in ± IFN-γ (100 U/ml for 24 h) activated WT and RNF213 KO A549 cells at 4 and 48 hpi. Data shows mean colony forming units (CFU) per well ± S.E.M. from four independent experiments each with 3 internal replicates. * *P* < 0.05 using two-tailed unpaired Student’s t-test. (B) Representative image showing co-localization of Mtb (green) with ubiquitin and RNF213. Cells were stained with FK2 (red) and RNF213 (magenta) in IFN-γ activated A549 cells that were analyzed at 8 hpi. Arrow depicts FK2+ and RNF213+ Mtb. Nuclei are labelled with Hoechst (blue). Scale bar = 5 μm. (C) Control and IFN-γ (24 h, 100 U/ml) activated WT, IRF1 KO and RNF213 KO A549 cells infected with MOI 1 VSV-GFP (recombinant vesicular stomatitis virus expressing enhanced green fluorescent protein) at 24 hpi. Bar plot shows levels of GFP expression plotted as mean arbitrary units (AU). Data are means ± SEM for 7 independent biological replicates. * *P* < 0.05 using Two-way ANOVA with Dunnett’s multiple comparison test. (D) Representative images showing total GFP emission imaged by Typhoon scanning of the 24-well plate.

## Discussion

To identify conserved mechanisms of cell autonomous immunity to intracellular pathogens, we performed a genome-wide CRISPR screen for loss of *T. gondii* growth restriction in response to IFN-γ. Although the primary screen failed to identify previous candidate genes implicated in control, we ascribed a new role for the tumor suppressor gene NF2 in upregulating expression of IRF-1 dependent ISGs. To account for the possibility that the primary screen lacked power to identify single gene effects, we performed an ISG-targeted screen for loss of IFN-γ-mediated *T. gondii* growth restriction. The ISG-targeted screen identified the E3 ligase RNF213 as the primary effector important for IFN-γ mediated control of *T. gondii* growth in human cells. RNF213 was upregulated by IFN-γ, recruited to the parasite containing vacuole, and associated with initial ubiquitination of targets at this interface. Deletion of RNF213 revealed that it was necessary for growth restriction, despite not relying on ATG5, which may act at downstream steps independent of growth restriction. Loss of RNF213 also diminished antibacterial and antiviral responses to interferon, suggesting it controls a common host pathway needed for pathogen resistance. These studies define a broad role for RNF213 as an essential mediator of growth restriction of divergent pathogens in a variety of human cell types.

Although previous studies have focused on candidate genes, we were interested in an unbiased method to identify genes important for IFN-γ mediated control of *T. gondii*. To this end, we developed a FACS-based CRISPR-screen to identify genes whose absence results in loss of IFN-γ mediated growth restriction of a GFP-expressing strain of *T. gondii*. The genome-wide CRISPR screen identified a number of components in the STAT1 signaling pathway including JAK2, the receptors IFNGR1 and IFNGR2, and the transcription factor IRF-1. These hits validate the screen as being sufficient to identify factors that globally affect the IFN-γ response. In particular, IRF-1 amplifies the responses normally triggered by engagement of STAT1 on promoters bearing GAS sequences (44), thus upregulating a constellation of ISGs. Consistent, with this ability we found that IRF-1 was necessary for maximal growth control following IFN-γ stimulation. Independently, we have previously shown that overexpression of IRF-1 in A549 cells is sufficient to induce control of *T. gondii* growth (34). We also identified NF2 as a significant hit in the primary genome-wide and pooled secondary CRISPR screens. Consistent with its prominent ranking in the CRISPR screens, deletion of NF2/Merlin significantly reduced the expression of IRF-1-dependent genes ungulated by IFN-γ, in addition to dysregulating numerous unrelated genes. This finding was unexpected as NF2 is known as a tumor suppressor gene that negatively regulates mTOR and the Hippo signaling pathways but is without prior connection to interferon signaling (40, 41). NF2/Merlin binds to and inhibits the E3 ubiquitin ligase CRL4 in the nucleus, thereby suppressing tumorigenesis (37). Previous studies have shown that CRL4 ubiquitinate histones and/or transcription factors leading to down-regulated gene expression (45). Hence, it is possible that in the absence of NF2/Merlin, CRL4 may target IRF1 to down-regulate its activity. Among the ISGs that are NF2 and IRF-1 dependent, RNF213 may factor prominently given its essential downstream role in mediating IFN-γ growth inhibition.

Because the genome-wide CRISPR screen failed to find previous candidate genes, we considered it might lack sufficient power to identify such single factors. As such, we performed a targeted CRISPR screen of ISGs that we have previously shown to be upregulated in A549 cells (34). Despite the fact that this screen included a number of previously recognized factors that control IFN-γ mediated growth in other human cells including RARESS3 (34), ISG15 (28), GBPs (32, 33), and IDO (25), these genes were not identified as significant hits. This finding suggests that while these genes have demonstrable effects when tested individually, their individual contributions are too modest to show differences at the population level, at least based on the conditions used here. Instead of these previously characterized factors, the ISG-targeted CRISPR screen identified a prominent role for RNF213, which is expressed at basal levels and upregulated in response to IFN-γ (34). RNF213 was essential for IFN-γ mediated restriction of *T. gondii* growth in multiple human cell types. We previously shown that type I IFN also restricts *T. gondii* growth in human THP-1 macrophages (3) and we now show that this restriction is downstream of RNF213.

We observed that RNF213 was recruited to a proportion of parasite containing vacuoles even in unstimulated cells, consistent with the fact that it is expressed at basal levels yet induced by IFN-γ. Similarly, RNF213 has been shown to be recruited to vacuoles containing intracellular *Salmonella* (42), *Listeria* (46), and *Chlamydia* lacking the effector GarD (47), and to restrict their growth in the absence of IFN-γ stimulation. We observed a slight increase in the growth of *T. gondii* in non-stimulated cells lacking RNF213; however, the major role for this factor was in IFN-γ treated cells which completely lost growth inhibition in the absence of RNF213. Notably, RNF213 was recruited early during infection and was associated with ubiquitination of targets on the parasite containing vacuole. It was also present there before autophagy marker LC3 and notably its presence on the vacuole did not rely on ATG5. Moreover, the growth restriction imparted by IFN-γ treatment was largely independent of ATG5 and yet fully dependent on RNF213. Taken together, these finding suggest a model whereby RNF213 is initially recruited to the parasite-containing vacuole where it ubiquitinates unknown targets leading to growth restriction, followed by recruitment of a non-canonical ATG pathway, which while not required for growth restriction may be important in downstream events.

The mechanism by which RNF213 is recruited to pathogen containing vacuoles remains undefined, but the broad specificity of this pathway suggests that there may be some common determinant recognized by the multiple domain structure of RNF213 that includes a dynein-like core with 6 ATPase units and a multi-domain E3 module (48). Similarly, the targets of ubiquitination on other pathogen-containing vacuoles remain undefined. Previous studies have suggested LPS on cytoplasmically exposed *Salmonella* is targeted by ubiquitination (42), although the exact chemical adduct remains to be defined. However, other pathogens including intracellular *Listeria*, and various viruses that have been shown to be targeted by RNF213 (46), lack LPS. The pathogens studied here also do not contain LPS, indicating that they must be recognized by a different mechanism. The broad nature of organisms targeted by RNF213 favors a model in which RNF213 recognizes a host target, either protein or lipid, whose modification leads to restricted pathogen growth. Future studies to elucidate this mechanism may enable enhanced cell-autonomous immunity to clear intracellular pathogens without the adverse effects of treatment with interferons.

## Acknowledgements

We thank Dr. Wandy Beatty, Microbiology Imaging Facility, Washington University in St. Louis for performing the electron microscopy studies. We also thank Jennifer Barks for cell culture support, members of the Sibley lab for helpful comments, Robert Orchard for advice on the CRISPR screens, and Felix Randow for communications prior to publication. This work was presented prior to publication at the Woods Hole Immunoparasitology Meeting (April 10-13, 2022) and the 16th International Congress on Toxoplasmosis (May 22-26, 2022). Supported in part by grants from the NIH (AI154048, AI118426).

## Author contributions

Conceptualization: SKM, LDS; Data curation, SKM; Investigation; SKM, HPK, PC; Formal Analysis, SKM, HPK, PC; Methodology: AB, JD; Project administration and supervision, JAP, SD, LDS; Visualization SKM, HPK, PC; Writing, SKM, SD, JAP, LDS.

## Database submission

RNASeq data have been submitted to GEO under the accession number GSE215771 and will be made available to reviewers and released prior to publication.

## Supplementary Materials

**Fig. S1A Validation of STAT1 knockout A549 line**.

**Fig. S1B Comparison of top hits for four primary CRISPR screens**.

**Fig. S2 Validation of IRF-1 and NF2 knockouts in A549 cells**.

**Fig. S3 Summary of NF2 dependent genes based on RNASeq**.

**Fig. S4 Validation of RNF213 knockouts in HFF and THP-1 cells**.

**Fig. S5 IFA staining of RNF213 and ubiquitination in CTG infected IFN-γ activated HFF and THP-1 cells**.

**Table S1 Primers used for sgRNA cloning**

**Dataset 1 Results of four primary CRISPR screens**.

**Dataset 2 Results of pooled secondary CRISPR screens**.

**Dataset 3 Results of ISG targeted CRISPR screens**.

## Materials and Methods

### Reagents and antibodies

Rabbit polyclonal anti-RNF213 (#HPA003347), Tryptophan, puromycin, polybrene and phorbol 12-myristate 13-acetate (PMA) were obtained from Sigma-Aldrich (St. Louis, MO, USA). Human IFN-γ, human TNF-α and human IFN-β were obtained from R&D systems (Minneapolis, MN, USA). *T. gondii* parasites were stained with mouse mAb DG52 against the surface antigen SAG1 (49) and GRA7 was detected using a rabbit polyclonal serum described previously (50). Mouse monoclonal anti-polyubiquitin (FK2, # 04-0263) antibody was obtained from Millipore Sigma Life Sciences. Rabbit mAb anti-IRF1 (D5E4, # 8478) and rabbit mAb anti-STAT1 (D1K9Y #14994) were obtained from Cell Signaling Technologies. Rabbit polyclonal LC3 (# PM036) was obtained from MBL International Corporation. Rabbit polyclonal anti-NF2 (ab217016) antibody was obtained from Abcam. Mouse monoclonal anti-ATG5 (#66744-1-Ig) was obtained from Proteintech Group, Inc. Secondary anti-IgGs conjugated to IRDye800 and IRDye700 were obtained from Li-Cor Biosciences (Lincoln, NE, USA). Hoechst, goat anti-mouse IgG and goat anti-rabbit IgG secondary antibodies conjugated to Alexa 488 or Alexa 594, were obtained from Life technologies Corporations (Grand Island, NY, USA). Secondary anti-IgG conjugated to 18 nm gold particle in immuno-electron microscopy was obtained from Jackson ImmunoResearch Laboratories, Inc., (West Grove, PA).

### Parasite and mammalian culture

Type III CTG expressing GFP parasites were generated previously (34). Parasites were grown as tachyzoites in human foreskin fibroblasts (HFF) obtained from the laboratory of Dr. John Boothroyd, Stanford University, as described previously (9). Parasites were harvested shortly after natural egress and purified by passage through 20 g needle and separated from host cell debris using 3.0 micron polycarbonate filters (Whatman). HFFs, HEK-293T and A549 lung epithelial carcinoma cells were maintained in DMEM with 10% v/v FBS at 5% CO_2_ and 37 °C. THP-1 human monocytic cells were maintained in RPMI with 10% v/v FBS at 5% CO_2_ and 37 °C. THP-1 cells were first differentiated into macrophages with 50 nM PMA for 48 h. PMA was washed off and cells were further incubated in their maintenance media for 24 h before use. All strains and host cell lines were determined to be mycoplasma-negative using the e-Myco Plus Kit (Intron Biotechnology).

### Lentiviral transduction of the Brunello library

The Brunello library (# 73179) expressing 76,441 sgRNAs against 19,114 human genes + 1,000 non-targeting sgRNA controls in plentiCRISPRv2 was obtained from Addgene (51). The library was electroporated into Endura electrocompetent cells (# 60242; Lucigen Bioresearch Technologies) and plasmid was prepared from > 80 million bacterial colonies (>1,000X representation). Briefly, twenty-five 60 mm tissue culture dishes, each containing 2.6 million 293T cells, were plated 24 h prior to transfection. A combination of 15 μg of plentiCRISPRv2 harboring the Brunello library, 5μg of pRSV-Rev (Addgene # 12253), 7 μg of pMD2.G (Addgene # 12259), and 11 μg of pMDLg/pRRE (Addgene # 12251) were co-transfected per tissue culture dish using Lipofectamine LTX reagent with PLUS reagent (Thermo Fisher Scientific) as per manufacturer’s instruction. The supernatant of 293T cells was collected 72 h after transfection and filtered using 0.45 μm PES filter. A total of ten 150 mm tissue culture dishes, each containing 10 million A549 cells, were seeded 24 h prior to virus transduction. The virus in the supernatant was serially diluted and transduced into A549 cells using polybrene (8 ug/ml) to achieve ∼ 10% transduction efficiency. The media was changed 72 h after transduction and cells were selected with 4 μg/ml of puromycin. In parallel, 800,000 A549 cells per well were seeded in 6-well plates to calculate the coverage of transduced Brunello library. Transduced coverage was calculated by measuring percent survival of virus transduced cells 72 h after incubation with 4 μg/ml puromycin relative to untreated control. The cells were selected on 4 μg/ml puromycin for two weeks to select the transduced cells. The Brunello library transduced A549 cells were passage 8 times before their use in the FACS based experiments. The library transduced cells were always spilt at a minimum of 100X coverage or 7.5 million cells in new T350 cm^2^ tissue culture flasks for every passage until completion of the experiment.

### Cloning of secondary sgRNA CRISPR library

Secondary screen: sgRNAs of genes that were significantly enriched (log2Fold Change > 1 and *P* < 0.05) in T5 over uninfected control in 2 out of 4 replicates of the primary Brunello library screens were selected to conduct a secondary screen. Ten sgRNAs per gene against 224 selected genes were designed using sgRNA designer at the Genetic Perturbation Platform (GPP) web portal (https://portals.broadinstitute.org/gpp/public/analysis-tools/sgrna-design). Additionally, 260 negative control sgRNAs were randomly selected from 1,000 negative control sgRNAs in the Brunello library with 130 each corresponding to NO_SITE and ONE_INTERGENIC_SITE negative controls. The oligonucleotide library was synthesized at Genescript. Briefly, the 20 bp sgRNAs in the library were flanked by complementary sequences in the plentiCRISPRv2 vector with CTTGTGGAAAGGACGAAACACCG oligonucleotide at the 5’ end and GTTTTAGAGCTAGAAATAGCAAGTTAAAATAAGGC at the 3’ end. The library was PCR amplified and Gibson cloned (52) into BsmBI digested plentiCRISPRv2. Assembled plasmid was dialyzed on a Type-VS Millipore membrane (MF type, VS filter, mean pore size = 0.025 μm, Millipore, Inc. #VSWP 02500) and electroporated into Endura electrocompetent cells. A549 cells were transduced with the secondary pool library at > 100X coverage, as described above, and selected with 4 μg/ml puromycin for two weeks. The secondary library transduced cells were passage 8 times before their use in the FACS based experiments and were always spilt at a minimum of 100X coverage for every passage until completion of the experiment.

### Cloning of the Interferon Stimulated Gene (ISG) sgRNA library

Genes that are significantly induced in A549 cells upon IFN-γ stimulation (34) were selected for sgRNA design. Ten sgRNA per gene were designed for 352 ISGs using the GPP web portal. A total of 480 negative control sgRNAs were randomly selected from 1,000 negative control sgRNAs from the Brunello library with 240 each corresponding to NO_SITE and ONE_INTERGENIC_SITE negative controls. The oligonucleotide library was synthesized at Genescript. The 20 bp sgRNAs in this library were flanked by BsmBI recognition sites along with overhang sequence for ligation with BsmBI digested plentiCRISPRv2. The 79 oligonucleotide library consists of 20 bp sgRNA sequences flanked with 5’ GCACTTGCTCGTACGACGCGTCTCACACCG sequence and 3’ GTTTCGAGACGTTAAGGTGCCGGGCCCAC sequence. The library was PCR amplified, digested with BsmBI, and ligated into plentiCRISPRv2 using NEB Golden Gate Assembly Kit (BsmBI-v2) (New England Biolabs Inc. # E1602S). The ligation reaction was dialyzed on a Type-VS Millipore membrane and electroporated into Endura electrocompetent cells. A549 cells were transduced with the ISG library at > 100X coverage, as described above and selected on 4 μg/ml puromycin for two weeks. The ISG library transduced cells were passage 8 times before their use in the FACS based experiments and were always spilt at a minimum of 100X coverage for every passage until completion of the experiment.

### Flow cytometry and cell sorting

Wild type and STAT1 knockout (STAT1 KO) A549 cells were cultured ± 100 U/ml IFN-γ for 24 h prior followed by washing with complete media. The cells were infected with CTG expressing GFP (CTG-GFP) at MOI 0.5 and cultured for 40 h. Cells were washed, trypsinized, and fixed with 4% paraformaldehyde followed by analysis on Sony SH800S cell sorter. Single cells were gated and GFP fluorescence was used to monitor intracellular growth of CTG-GFP. FACS-based CRISPR screens were executed by pre-activating 30-50 million library transduced A549 cells with 100 U/ml of IFN-γ for 24 h. The cells were washed and infected with CTG-GFP at MOI 0.5 and incubated for 40 h. Cells were washed, trypsinized, and filtered (40 μm) prior to sorting. Single cells were gated and top 5% of events (T5) based on GFP fluorescence intensity were sorted. Genomic DNA was isolated from the top 5% CTG-GFP infected cells and uninfected cells for each of 4 biological replicates for each library. Uninfected control cells for each replicate were passage matched with the sorted T5 population.

### Genomic DNA isolation and next generation sequencing

Genomic DNA was isolated using Qiagen DNeasy kit as per manufacturer’s instructions. PCR was done using Ex Taq in a 100 μl reaction with the maximum of 10 μg of template per reaction (36). The amplicons for next generation sequencing were generated using a 2-step PCR amplification of integrated sgRNAs. Briefly, genomic DNA isolated from different samples was used as template for PCR1 using forward 5’ TCGTCGGCAGCGTCAGATGTGTATAAGAGACAGGGCTTTATATATCTTGTGGAAAGGACGA AACACC 3’ and reverse 5’ GTCTCGTGGGCTCGGAGATGTGTATAAGAGACAGCCAATTCCCACTCCTTTCAAGACCT 3’ primers with PCR cycling conditions of 30s 95 °C, 30s 53 °C, 30s 72 °C for 18-24 cycles. PCR2 was performed using 5 μl of PCR1 as template using forward 5’ AATGATACGGCGACCACCGAGATCTACAC-10bp barcode-TCGTCGGCAGCGTC 3’ and reverse 5’ CAAGCAGAAGACGGCATACGAGAT-10bp barcode-GTCTCGTGGGCTCGG 3’ primers with PCR cycling conditions of 30s 95 °C, 30s 54 °C, 30s 72 °C for 24 cycles. PCR2 amplicons were uniquely dual indexed with 10 bp Illumina-compatible barcodes for each sample. The samples within each screen were pooled and submitted to the Genome Technology Access Center, Washington University School of Medicine in St. Louis for next generation sequencing on an Illumina NovaSeq-6000 with at least 100x coverage of 150 bp paired end reads. Basecalls and demultiplexing were performed with Illumina’s bcl2fastq2 software. The forward reads were analyzed using MAGeCK as described previously (53). Briefly, read counts for the sgRNAs sequences were normalized and counted using the count function followed by comparison between the T5 population and uninfected control using the test function. The data was analyzed in R using the MAGeCKFlute package (54). Log_2_Fold change of sgRNAs in T5 population relative to uninfected control was plotted against -Log_10_FDR (False Discovery Rate).

### RNA Sequencing

Cells were seeded in a 6-well plate overnight and then treated ± IFN-γ (100 U/ml) for 24 h. The cells were lysed, and total RNA was extracted using QIAGEN RNeasy Mini Kit as per manufacturer’s instructions. The RNA samples were prepared from 3 independent biological replicates and submitted to the Genome Technology Access Center, Washington University School of Medicine in St. Louis, for next-generation mRNA sequencing. Library preparation was performed using ∼ 1ug of total RNA processed using RiboErase kits to remove ribosomal RNA (Kapa Biosystems). mRNA was then fragmented in reverse transcriptase buffer and heating to 94 °C for 8 min. mRNA was reverse transcribed to yield cDNA using SuperScript III RT enzyme (Life Technologies, per manufacturer’s instructions) using random hexamers. A second strand reaction was performed to yield ds-cDNA that was blunt ended, extended by adding an A base to the 3’ ends, and followed by ligation of Illumina sequencing adapters. Ligated fragments were then amplified for 12-15 cycles using primers incorporating unique dual index tags. Fragments were sequenced on an Illumina NovaSeq-6000 to generate paired end reads extending 150 bases. Basecalls and demultiplexing were performed with Illumina’s bcl2fastq2 software. A total of 1,046,709,676 reads from 18 samples were generated and the fastq files were imported into CLC Genomics Workbench version 21.0.5 (QIAGEN Bioinformatics, Inc.). The reads were aligned to Homo sapiens hg38 reference genome (downloaded from Ensembl via the CLC Genomics Workbench) that covered 60,676 genes and 204,381 transcripts in total. The expression data from the CLC Genomics Workbench were imported into R to generate heat map of differentially expressed genes and compare RPKM values of ISGs across samples.

### CRISPR-Cas9 mediated gene deletions

sgRNAs were designed using the GPP web portal and 20 bp sgRNAs were synthesized as primer pairs (**Table S1.xlsx**) with overhanging BsmBI sites and ligated to BsmBI digested plentiCRISPRv2. sgRNAs cloned into the plentiCRIPSRv2 plasmid was packaged into lentiviruses by co-transfecting into 293T cells in equimolar ratio with pRSV-Rev, pMD2.G and pMDLg/pRRE. The supernatant was collected 72 h post transfection and the virus was filtered through 0.45 μm PES filter. The virus containing supernatant was titrated and was transduced into different cell lines at 10% transduction efficiency for 72 h. The cells were washed and incubated with puromycin (4 μg/ml for A549, 2 μg/ml for THP-1 and 1 μg/ml HFFs) for 10 days to select stably transduced cells. Single cell knockouts of IRF-1, NF2, and RNF213 were isolated by FACS sorting single cell per well. STAT1 KO A549 cells, ATG5 KO A549 cells and RNF213 KO HFF cells were maintained at the population level. Gene deletion of host factors was confirmed by western blot and Sanger sequencing of the editing site.

### Vacuolar size growth assay

Parasite growth was monitored using a vacuolar size assay that was performed as described previously (9). Briefly, host cells were seeded in 96-well μCLEAR black plates (Greiner Bio-One International GmbH) for 24 h before use. For THP-1, the cells were first differentiated with 50 nM PMA for 48 h before addition of 100 U/ml of IFN-γ or 100 U/ml IFN-β with 10 ng/ml of TNF-α for 24 h prior to infection. The cells were cultured ± 100 U/ml of IFN-γ for 24 h and then washed with warm PBS. The cells were infected with parasites at MOI 0.5 for 2 h, washed with warm PBS to remove extracellular parasites, and returned to culture in complete medium for different intervals. Cells were fixed with 4% paraformaldehyde, permeabilized using 0.02% saponin in 5% FBS (Fetal bovine serum) +5% NGS (Normal goat serum) and stained to detect parasites (anti-SAG1) and the parasitophorous vacuole (anti-GRA7), followed by Alexa-fluor conjugated secondary antibodies, and Hoechst (100 ng/ml) to stain the nuclei. Images were acquired at a magnification of 20x on a Cytation3 cell imaging multi-mode plate-based imager (BioTek). The size of parasitophorous vacuole-harboring parasites (SAG1-positive GRA7 vacuoles) was determined using CellProfiler 2.1.1. Data from at least 50 fields per sample in each of three independent experiments were used to calculate the vacuolar size of parasites in different host cells.

### ISRE promoter assay

Cells per well were seeded at a density of 25,000 per 96-well plate and cultured overnight. The cells were co-transfected with 50 ng of plasmid expressing ISRE reporter (9) and 50 ng of pCMV-Red Firefly Luc Vector (Thermo Fisher Scientific # 16156) using lipofectamine LTX (Thermo Fisher Scientific) according to the manufacturer’s instructions. The cells were washed with warm DMEM (with 10% v/v FBS) and cultured ± 100 U/ml human IFN-β for another 24 h. Gaussia luciferase activity was measured in the supernatant using the Pierce Gaussia luciferase glow assay kit (Thermo Fisher Scientific # 16160). Firefly luciferase activity was measured in the cell lysates after removing the supernatant and washing the cells with warm PBS using the Pierce Firefly luciferase glow assay kit (Thermo Fisher Scientific # 16176). Luminescence was measured in Cytation3 multi-mode plate reader. ISRE promoter activity was calculated by normalizing gaussian luminescence to firefly luminescence for each sample.

### Immunofluorescence staining

Cells were seeded onto 12 mm coverslips in a 24-well overnight and then treated ± 100 U/ml human IFN-γ for 24 h. The cells were washed with warm PBS and infected with CTG-GFP parasites at MOI 3 for 6h. The cells were washed to remove extracellular parasites and fixed with 4% paraformaldehyde for 20 min at room temperature. Cells were permeabilized and blocked in using 5% FBS and 5% normal goat serum with 0.02% saponin in PBS for 30 min. Cell were stained with FK2 (1:2,000; anti-mono- and poly-ubiquitinated proteins) and anti-RNF213 (1:2,000) and/or anti-LC3 (1:2,000) in 1% FBS with 0.02% saponin in PBS for 90 min. Cells were washed three times with PBS and incubated with Alexa-fluor conjugated secondary antibodies (1:4,000) and Hoechst stain (100 ng/ml) to stain nuclei (Life Technologies) for 30 min. Cells were then washed three times with PBS followed by mounting in ProLong gold antifade mountant (Thermo Fisher Scientific). Images were acquired using a 63x oil Plan-Apochromat lens (N.A. 1.4) on an Axioskop2 MOT Plus Wide Field Fluorescence Microscope (Carl Zeiss) or using a LSM880 confocal laser scanning microscope (Carl Zeiss) in the Microbiology Imaging Facility, Washington University in St. Louis.

### Western blotting

Cells were lysed using CellLytic M (Sigma) with complete mini protease inhibitor cocktail (Roche). The lysates were centrifuged at 6,000 *g* for 10 min at room temperature to pellet nuclear debris. Total protein was measured using BCA (bicinchoninic acid) protein assay kit (Thermo Fisher Scientific). Samples were boiled at 95 °C for 15 min in Laemmli buffer containing 100 mM dithiothreitol (DTT), separated by SDS-PAGE and transferred onto nitrocellulose membranes. Membranes were blocked in a 1:1 mixture of Odyssey blocking buffer (OBB; Li-Cor Biosciences) and PBS overnight at 4°C. Membranes were incubated with primary antibodies (1:4,000) for 2 h at room temperature in a 1:1 mixture of Odyssey blocking buffer and PBS with 0.1% Tween 20 (PBST). Blots were washed thrice with PBST and incubated with anti-mouse IgG IR800 and anti-mouse IgG IR700 at 1:15,000 for 2 h at RT in a 1:1 mixture of Odyssey blocking buffer and PBST. The blot was washed thrice with PBST followed by infrared imaging on a Li-Cor Odyssey imaging system.

### *M. tuberculosis* strains and infections

*M. tuberculosis* (Mtb) was grown at 37 °C to log phase in Middlebrook 7H9 broth (BD Biosciences) supplemented with 0.05% Tyloxapol, BD BBL Middlebrook OADC Enrichment (BD Biosciences), and 0.2% (v/v) glycerol. H37Rv (the wild type strain) and H37Rv-GFP (GFP-expressing fluorescent strain) were originally provided by William Jacobs Jr. (Albert Einstein College of Medicine). H37Rv-GFP was maintained in broth containing 25 μg/mL kanamycin. For *in vitro* infection assays, a log phase culture of *M. tuberculosis* was used to prepare single-cell suspensions using the slow centrifugation method. Bacteria were pelleted, resuspended in cell culture medium, and centrifuged at 800 rpm for 8 minutes. The supernatant was collected, and the number of bacilli was estimated by measuring absorbance at 600 nm. This step was repeated until a constant absorbance value was obtained. For *in vitro* infection, cells were seeded in 96 well plates overnight followed by culture ± human IFN-γ (100 units/mL) for 24 h. Cells were washed and infected with Mtb at MOI 0.5. After 4 hours, cells were washed three times with warm DMEM to remove extracellular bacteria, and then resuspended in culture medium. To estimate intracellular bacterial growth, infected cells were lysed in 0.06% SDS solution at the indicated time points, and serial dilutions of the lysates were plated on 7H11 agar plates (BD Biosciences, catalog no. 283810) containing BD BBL Middlebrook OADC Enrichment (BD Biosciences, catalog no. 212351) and glycerol. Colony forming units (CFU) were calculated 14-21 days later. For immunofluorescence assays, H37Rv-GFP infected cells were fixed at 8 hpi with 4% paraformaldehyde for one hour, permeabilized and blocked in PBS with 0.05% Triton X-100 and 3% BSA and stained with FK2 (1:2,000) and anti-RNF213 (1:2,000) overnight at 4 °C. Staining with Alexa fluorophore-conjugated secondary antibody (Molecular Probes) was done for 2 h at room temperature. Following this, the samples were washed with 0.1% Tween 20/PBS and mounted using Prolong Gold antifade (Thermo Fisher Scientific, # P36930). Images were captured using a Nikon Eclipse Ti confocal microscope (Nikon Instruments Inc.) equipped with a 60X apochromat oil-objective lens and analyzed using NIS-Elements version 4.40 (Nikon).

### Vesicular Stomatitis virus (VSV) culture and infections

Recombinant vesicular stomatitis virus (VSV) expressing enhanced green fluorescent protein (eGFP; VSV-GFP) (55) was propagated as previously described (56) in either African Green Monkey kidney Vero E6 (CRL-1586, ATCC) or MA104 cells (CRL-2378.1). Vero E6 cells and MA104 cells were cultured in complete DMEM media and complete M199 media, respectively. Viral titers were determined by standard plaque assays. Cells were seeded at 10,000 per well in 24-well plates and cultured overnight followed by culture in ± 100 U/ml human IFN-γ for 24 h. The cells were washed and infected with VSV-GFP at MOI 1 for 24 h, as described previously (57). GFP intensity was determined by scanning with an Amersham Typhoon 5 (GE) and images quantified by ImageJ. The number of total intensities subtracted from background was computed for each treatment and log_10_ transformed.

### Immuno-electron microscopy

Wild type A549 cells were pre-activated with IFN-γ (100 U/ml) 24 h prior to infection with CTG for 6h. The cells were trypsinized and fixed in 4% paraformaldehyde/0.01% glutaraldehyde (Polysciences Inc., Warrington, PA) in 100 mM PIPES/0.5 mM MgCl_2_, pH 7.2 for 1 h at 4 °C. Samples were then embedded in 10% gelatin and infiltrated overnight with 2.3 M sucrose/20% polyvinyl pyrrolidone in PIPES/MgCl_2_ at 4 °C. Samples were trimmed, frozen in liquid nitrogen, and sectioned with a Leica Ultracut UCT7 cryo-ultramicrotome (Leica Microsystems Inc., Bannockburn, IL). Ultrathin sections of 50 nm were blocked with 5% FBS (fetal bovine serum)/5% NGS (normal goat serum) for 30 min and subsequently incubated with rabbit anti-RNF213 antibody for 1 h at room temperature. Following washes in block buffer, sections were incubated with secondary anti-rabbit IgG antibody conjugated to 18 nm colloidal gold (Jackson ImmunoResearch Laboratories, Inc., West Grove, PA) for 1 h. Sections were stained with 0.3% uranyl acetate/2% methyl cellulose and viewed by transmission electron microscopy. All labeling experiments were conducted in parallel with controls omitting the primary antibody.

### Transmission electron microscopy

Wild type A549 cells were pre-activated with IFN-γ (100 U/ml) 24 h prior to infection with CTG for 6h. The cells were trypsinized and fixed freshly prepared mixture of 1% glutaraldehyde (Polysciences Inc, Warrington, PA) and 1% osmium tetroxide (Polysciences Inc.) in 50 mM phosphate buffer at 4ºC for 30 min. Samples were then rinsed extensively in cold dH_2_0 prior to en bloc staining with 1% aqueous uranyl acetate (Ted Pella Inc., Redding, CA) at 4 ºC for 3 h. Following several rinses in dH_2_0, samples were dehydrated in a graded series of ethanol and embedded in Eponate 12 resin (Ted Pella Inc.). Sections of 95 nm were cut with a Leica Ultracut UCT ultramicrotome (Leica Microsystems Inc., Bannockburn, IL), stained with uranyl acetate and lead citrate, and viewed on a JEOL 1200 EX transmission electron microscope (JEOL USA Inc., Peabody, MA) equipped with an AMT 8 megapixel digital camera and AMT Image Capture Engine V602 software (Advanced Microscopy Techniques, Woburn, MA).

### Statistical analysis

All experiments were performed in 3 independent biological repeats unless mentioned otherwise. Statistical analysis for each experiment was performed on the combined data in PRISM. Significance values and tests performed are included in the figure legends.

