## Supplemental materials for "Molecular Basis for Interferon-mediated Pathogen Restriction in Human Cells" (from another version of this preprint: version 1, 2022-10-21)

Supplementary Material

A

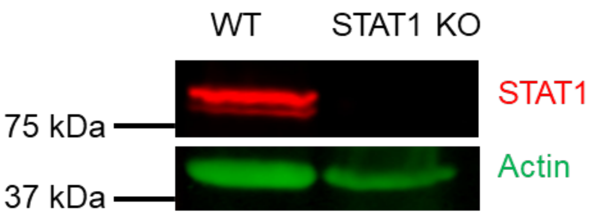


B

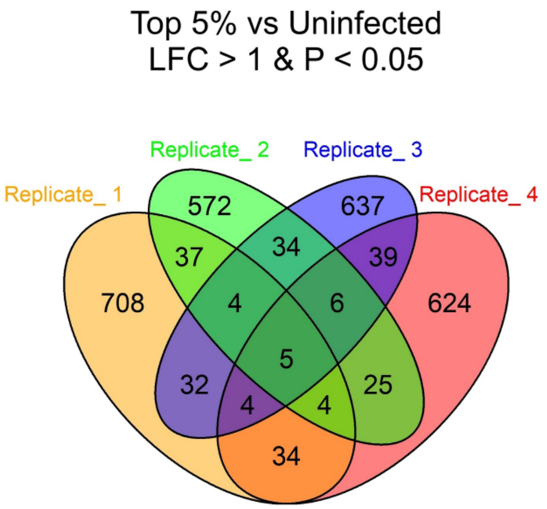


Genes enriched in

4/4 replicates: 5 (NF2, FRYL, UBE2A, CCDC6, PDCD10)

3/4 replicates: 23

2/4 replicates: 224

**Fig. S1** (A) Cellular lysates of wild type and STAT1 KO A549 cells were immunoblotted with anti-STAT1 antibody. Actin was used as loading control. (B) Venn diagram of genes significantly enriched with Log2 Fold Change > 1 and *P* < 0.05 (Raw *P* -value using permutation) in top 5% (T5) of the FACS sorted population compared to uninfected controls for each of the four replicates of primary Brunello screen.

A

```
>IRF1_Clone4
Sequence ID: WT_59609 Length: 453
Range 1: 1 to 453

Score:819 bits(443), Expect:0.0,
Identities:453/457(99%),  Gaps:4/457(0%), Strand: Plus/Plus
                                                                        sgRNA target
WT       181   TTCATGGCACAGCGAAAGTTGGCCTTCCACGTCTTGGGATCTGGCTCCTTTTCCCCTGCT   240
               ||||||||||||||||||||||||||||||||    ||||||||||||||||||||||||
IRF1 KO  181   TTCATGGCACAGCGAAAGTTGGCCTTCCACGT----GGATCTGGCTCCTTTTCCCCTGCT   236
```

B

```
>NF2_Clone7
Sequence ID: WT_26069 Length: 198
Range 1: 1 to 198

Score:359 bits(194), Expect:2e-104,
Identities:197/198(99%),  Gaps:1/198(0%), Strand: Plus/Plus
                                                  sgRNA target
WT      1    AAGGGAAGGACCTCTT-TGATTTGGTGTGCCGGACTCTGGGGCTCCGAGAAACCTGGTTC   59
             |||||||||||||||| |||||||||||||||||||||||||||||||||||||||||||
NF2 KO  1    AAGGGAAGGACCTCTTATGATTTGGTGTGCCGGACTCTGGGGCTCCGAGAAACCTGGTTC   60
```

**Fig. S2** Confirmation of single cell knockouts using Sanger sequencing. Basic local alignment of nucleotide sequence of sgRNA targeted site in the genome of (A) IRF1 KO and (B) NF2 KO A549 cells compared to wild type.

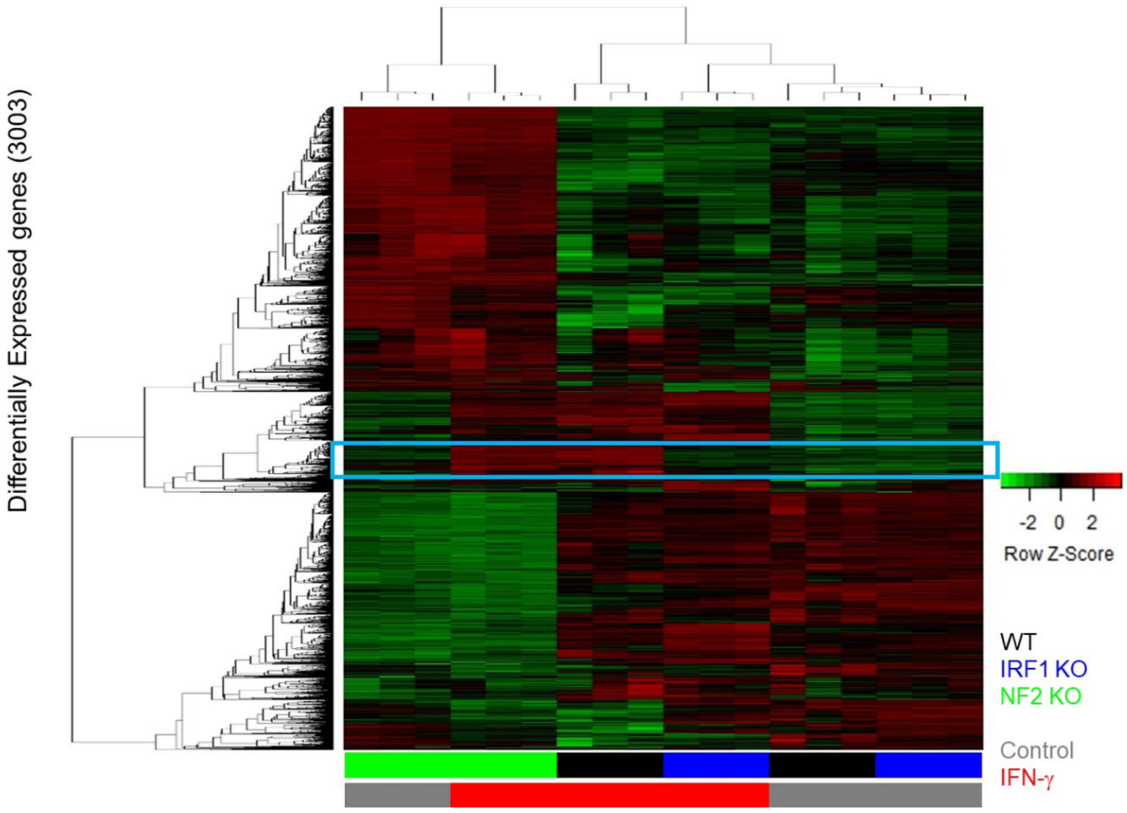


**Fig. S3** Heatmap of gene expression of significantly expressed genes (-1 ≤ Log2 Fold Change ≥ 1 and *P* < 0.05) across all samples with their 3 independent biological replicates clustered using 1 - Pearson distance and average linkage on normalized $Log_2$ (RPKM) values. Normalized Z-scores are color-scaled from green to red showing relative downregulation to upregulation. Genes that were induced upon IFN-γ activation in wild type cells but not in IRF1 KO cells are highlighted in cyan colored box.

A

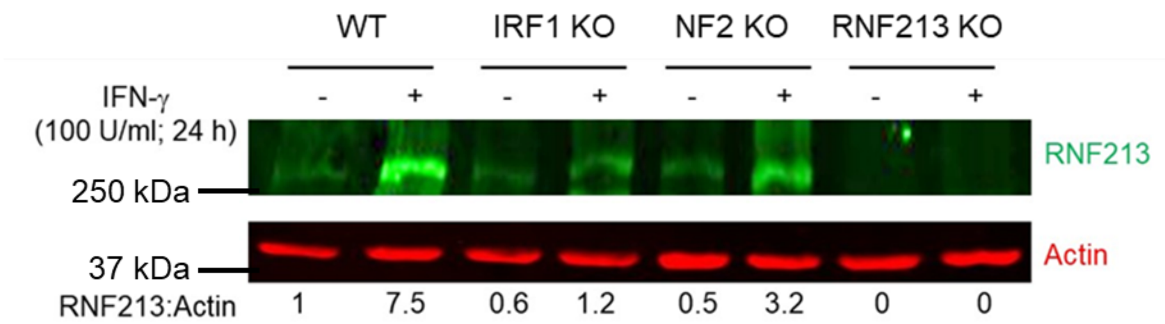


B

```
>RNF213_CloneC6 Allele 1
Sequence ID: WT_43109 Length: 240
Range 1: 1 to 240

Score:438 bits(237), Expect:4e-128,
Identities:240/241(99%),  Gaps:1/241(0%), Strand: Plus/Plus
                                                          sgRNA target
WT          61   CCCGTGCTTGTTCCCGGGCTCAGACAGTTGGCAAGAAAACCCCGAGGAGCCCTGTTCCAA  120
                 ||||||||||||||||||||||||||||||||||||||| ||||||||||||||||||||
RNF213 KO   61   CCCGTGCTTGTTCCCGGGCTCAGACAGTTGGCAAGAAAA-CCCGAGGAGCCCTGTTCCAA  119

>RNF213_CloneC6 Allele 2
Sequence ID: WT_34359 Length: 246
Range 1: 1 to 246

Score:399 bits(216), Expect:2e-116,
Identities:237/246(96%),  Gaps:5/246(2%), Strand: Plus/Plus
                                                                sgRNA target
WT          61   CCCGTGCTTGTTCCCGGGCTCAGACAGTTGGCAAGAAAACC--CC-GAGG-A-GCCCTGT  115
                 ||||||||||||||||||||||||||||||||||||||||   || |    | |||||||
RNF213 KO   61   CCCGTGCTTGTTCCCGGGCTCAGACAGTTGGCAAGAAAACTGTCCTGTTCCAGGCCCTGT  120
```

C

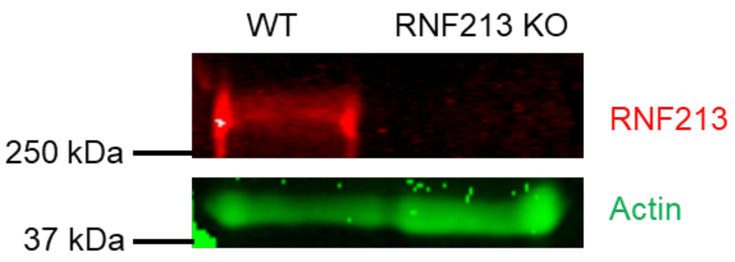


D

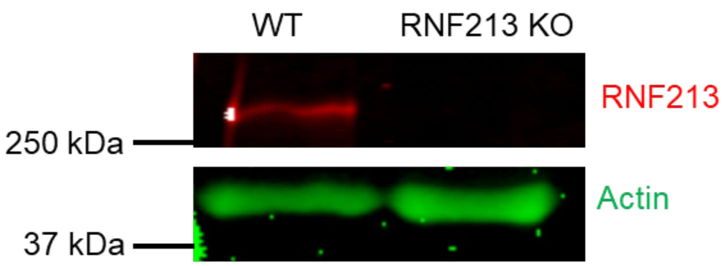


E

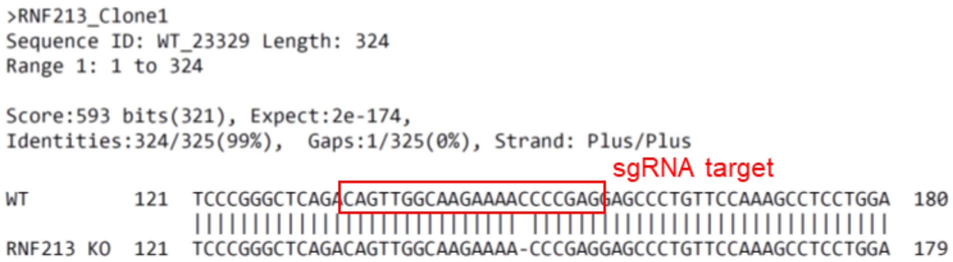


**Fig. S4** Cell lysates of wild type, IRF1 KO, NF2 KO and RNF213 KO A549 cells were immunoblotted using anti-RNF213 antibody. Actin was used as loading control. RNF213 specific band intensity was normalized to actin for comparison across samples. (B) Confirmation of RNF213 KO single cell knockout using Sanger sequencing. Alignment of the nucleotide sequence of sgRNA targeted site in the genome of RNF213 KO A549 cells compared to wild type. The 2 alleles of the single knockout were deconvolved using CRISPR-ID (http://crispid.gbiomed.kuleuven.be/). (C) Cell lysates of wild type and RNF213 KO HFF cells immunoblotted using anti-RNF213 antibody. Actin was used as loading control. (D) Cell lysates of wild type and RNF213 KO THP-1 cells were immunoblotted using anti-RNF213 antibody. Actin was used as loading control. (E) Alignment of the nucleotide sequence of sgRNA targeted site in the genome of RNF213 KO THP-1 cells compared to wild type.

A

HFFs

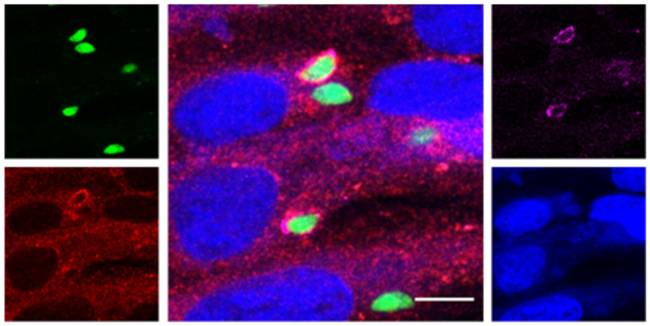


B

THP-1

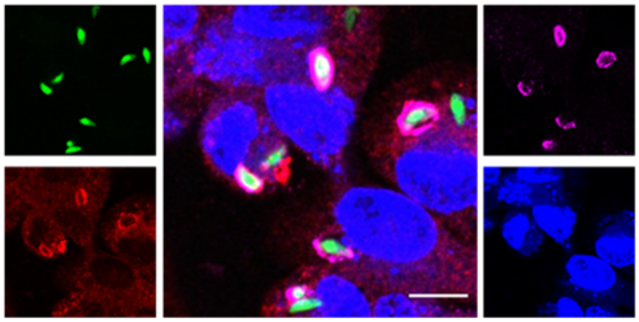


**Fig. S5** IFN-γ activated wild type and RNF213 KO (A) HFFs and (B) THP-1 cells infected with CTG-GFP at 6 hpi. The cells were fixed and stained with FK2 (red) and anti-RNF213 (magenta). Nuclei were labelled with Hoechst (blue) and parasites were GFP labelled (green), Bar = 10 μm.

**Table S1 Primers used for sgRNA cloning**

| Gene | Exon | Position | Forward | Reverse |
|---|---|---|---|---|
| STAT1 | 18 | 1543 | CACCGCACCAAAAGAGGTCTCAATG | AAACCATTGAGACCTCTTTTGGTGC |
| IRF1 | 4 | 225 | CACCGAGGAGCCAGATCCCAAGACG | AAACCGTCTTGGGATCTGGCTCCTC |
| NF2 | 2 | 140 | CACCGTCCGGCACACCAAATCAAAG | AAACCTTTGATTTGGTGTGCCGGAC |
| RNF213 | 3 | 220 | CACCGCAGTTGGCAAGAAAACCCCG | AAACCGGGGTTTTCTTGCCAACTGC |
| ATG5 | 2 | 54 | CACCGTGATATAGCGTGAAACAAGT | AAACACTTGTTTCACGCTATATCAC |
